# Dimensional Reduction for Single Molecule Imaging of DNA and Nucleosome Condensation by Polyamines, HP1α and Ki-67

**DOI:** 10.1101/2023.01.04.522433

**Authors:** Nils A. Benning, Jacob Kæstel-Hansen, Fahad Rashid, Sangwoo Park, Raquel Merino Urteaga, Ting-Wei Liao, Jingzhou Hao, James M. Berger, Nikos S. Hatzakis, Taekjip Ha

**Affiliations:** Department of Biology, Johns Hopkins University, Baltimore, MD 21218, USA; Department of Chemistry and Nanoscience Centre, University of Copenhagen, Copenhagen, Denmark; Department of Biophysics and Biophysical Chemistry, Johns Hopkins University School of Medicine, Baltimore, MD, 21205, USA; Department of Biophysics, Johns Hopkins University, Baltimore, MD 21218, USA; Novo Nordisk Foundation Centre for Protein Research, Faculty of Health and Medical Sciences, University of Copenhagen, Copenhagen, Denmark; Department of Biomedical Engineering, Johns Hopkins University, Baltimore, MD 21218, USA; Howard Hughes Medical Institute, Baltimore, MD 21205, USA

**Keywords:** Supported Lipid Bilayer (SLB), Spermine, HP1α, Ki-67, Particle Tracking, Diffusion, Nucleosome

## Abstract

Macromolecules organize themselves into discrete membrane-less compartments. Mounting evidence has suggested that nucleosomes as well as DNA itself can undergo clustering or condensation to regulate genomic activity. Current in vitro condensation studies provide insight into the physical properties of condensates, such as surface tension and diffusion. However, such studies lack the resolution needed for complex kinetic studies of multicomponent condensation. Here, we use a supported lipid bilayer platform in tandem with total internal reflection microscopy to observe the 2-dimensional movement of DNA and nucleosomes at the single-molecule resolution. This dimensional reduction from 3-dimensional studies allows us to observe the initial condensation events and dissolution of these early condensates in the presence of physiological condensing agents. Using polyamines, we observed that the initial condensation happens on a timescale of minutes while dissolution occurs within seconds upon charge inversion. Polyamine valency, DNA length and GC content affect threshold polyamine concentration for condensation. Protein-based nucleosome condensing agents, HP1α and Ki-67, have much lower threshold concentration for condensation than charge-based condensing agents, with Ki-67 being the most effective as low as 100 pM for nucleosome condensation. In addition, we did not observe condensate dissolution even at the highest concentrations of HP1α and Ki-67 tested. We also introduce a two-color imaging scheme where nucleosomes of high density labeled in one color is used to demarcate condensate boundaries and identical nucleosomes of another color at low density can be tracked relative to the boundaries after Ki-67 mediated condensation. Our platform should enable the ultimate resolution of single molecules in condensation dynamics studies of chromatin components under defined physicochemical conditions.

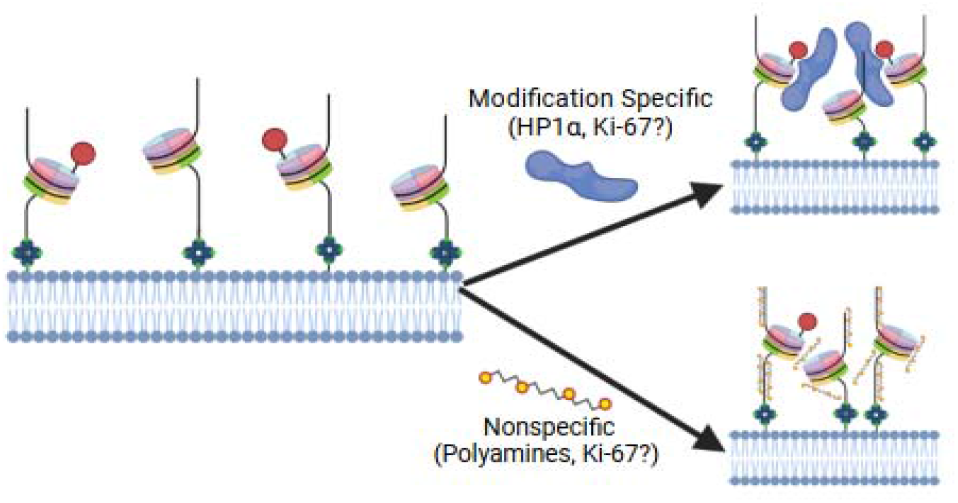

## 1. INTRODUCTION

A resurgent concept in biology is the way cells organize bio-molecules into membrane-less compartments through condensation. This organization facilitates cellular reactions and occurs once their concentration reaches a certain threshold. In the nucleus, these condensates facilitate processes such as DNA damage repair, gene repression, and ribosome biogenesis^2–4^; and in the cytosol they facilitate mRNA processing, mRNA localization, translation, protein folding, along with many other processes^5^. Dysregulation of condensates may result in the formation of pathological aggregates that lead to impaired cell function and may ultimately trigger cell death^5–15^.

Biomolecular condensation is promoted through interactions between macromolecules and depends on valency of interactions. Entropy typically favors a heterogenous mixture and is governed by environmental factors like pH and temperature, but many biological condensates circumvent this through multivalent interactions. Many proteins achieve multivalency through the interaction of two generally conserved modules: folded domains and low complexity disordered segments^5,2,16^. While recent advances detail the types of multivalent interactions and potential critical concentrations required for condensation, many studies lack the resolution required to distinguish phase-separated condensates from the formation of complex ordered structures ^17,18^. In addition, while live-cell based studies have examined protein constructs with tunable valency^19,20^, the kinetics of early condensate formation remain mostly unexplored.

Single molecule studies provide enhanced resolution to observe the movement of individual macromolecules and how these mobile particles interact^21, 22^. Here, we used a supported lipid bilayer (SLB) to investigate 2-dimensional movement of DNA and nucleosomes early in their condensation at the single-molecule level. Previously, SLBs have been used to study vesicle fusion, cell adhesion^23–29^, and importantly, the clustering of components within the bilayer like individual lipids and cholesterol^30^. This platform allows us to correlate kinetic behavior with condensate properties of SLB-bound molecules. Thus, it becomes possible to investigate the recruitment kinetics of other phase separating proteins or macromolecules as we show for polyamines and HP1α. One protein of particular interest is Ki-67, a 2896 amino acid protein that helps maintain chromosome individuality by coating nucleosomes during mitosis^37^. At the end of mitosis, Ki-67 aids in the exclusion of cytoplasm during the reformation of the nuclear envelope. Thus, we used our experimental platform to examine Ki-67 as a condensing agent for DNA and nucleosomes.

## 2. METHODS

### 2.1 Supported lipid bilayer (SLB) generation

SLBs were generated based on studies that observe vesicle fusion into bilayers^25^. Small unilamellar vesicles are prepared by drying a mixture of 93% POPC and 7% 18:1 biotinyl-PE (Avanti Polar Lipids, catalog #850457C and 870282C) under compressed nitrogen gas followed by overnight drying under vacuum. This “lipid cake” was hydrated with T50 Buffer (10 mM Tris-HCL pH 8.0, 50 mM NaCl) and pipetted several times to promote vesicle formation. After undergoing 15 freeze-thaw cycles using liquid nitrogen, small unilamellar vesicles (SUVs) were prepared by 21 passes through an extruder (Avanti) fitted with 100 nm filter (Cytiva, catalog # 800309). SUVs were stored at 4°C for up to 14 days.

### 2.2 Slide Preparation/Assembly

Quartz slides were cleaned by sonicating in methanol for 30 min, then washing with acetone and drying with nitrogen gas. After drying, the slides were sonicated in a 5% Alconox detergent solution for 30 min and rinsed with ddH_2_O. Slides were then sonicated in a 1 M KOH solution, rinsed with ddH2O, and burned using a propane torch. Glass coverslips were incubated in a 1X detergent solution (MP Biomedicals, 097667093) just below boiling point for 1 hour and then thoroughly rinsed with MilliQ water. Glass coverslips were then baked in a furnace (Barnstead International, Model FB1315M) at 540 °C, just below the melting point of the coverslip, for 5 hours. Once clean, 1-2 mm strips of Scotch double-sided tape were placed between pre-drilled holes on the edges of the quartz slide. The glass coverslips were then briefly burned using a propane torch and placed on the tape covered quartz slide. Excess tape was removed, and the edges of the slide were sealed using epoxy (Devcon), resulting in a final assembled reaction chamber. Functional imaging slides are made by injecting 20 μL of SUVs into a reaction chamber and incubating for 30 minutes to promote vesicle fusion and SLB formation. The excess free SUVs were then washed out with 200 μL of T50 Buffer.

### 2.3 TIRF microscopy/ Analysis

Fluorescently labeled macromolecules (DNA and mono-nucleosomes) were observed on the SLBs using objective-based total internal reflection fluorescence microscopy. The fluorescence emission was collected by oil objective (Nikon PlanApo, NA 1.40, 60×) and recorded by a back-illuminated electron-multiplying charge-coupled device camera (iXon3, Andor Technology) with 50 ms exposure time.

To measure initial condensation events, two criteria were put in place to define a potential condensate: no diffusion, and an intensity value at least double that of an individual particle. Once a potential condensate is identified, other potential condensates can be easily located with the same criteria. When potential condensates are visually identified after moving the field of view to a random position or multiple condensates appear within one field of view, initial condensation has occurred.

Single molecule tracking and fluorescence intensity analysis was performed using the ImageJ plugin, TrackMate, with a Laplacian of Gaussian (LoG) filter detector, which enables analysis of merging and splitting events. XY coordinates were obtained from these videos for each particle in each frame, and trajectories for each particle were analyzed using a Linear Assignment Problem (LAP) tracker^31^. Diffusion coefficients for each particle were determined using a mean squared displacement (MSD) analysis. Particles were assumed to undergo simple diffusion (where MSD = 4Dt), i.e.: no confinement or directed movement. Trajectories were excluded when trajectory length was fewer than 10 frames, indicating particles transiently entered the field of view. To track real-time diffusional behavior, we utilized rolling MSD analysis by fitting MSD = 2dDt^α^, where D is the diffusional coefficient, α is a measure of persistence in the walk and d is the dimension. For the analysis we applied a system-specific threshold for restricted motion of MSD=0.05 μm^2^.

### 2.4 Widefield microscopy/ Analysis

Bulk diffusion measurements were obtained by preparing SLBs as described above and imaging via a 555 nm excitation laser with a Cy3 emission filter using a water/oil-immersion objective (Nikon PlanApo λ, NA 0.75, 20x). FRAP experiments were performed using this microscope with a 50 mW bleaching laser at 405 nm and a Bruker Galvano mirror scanner. Regions of interest measuring 25 pixels in diameter on the SLB were bleached using 50% laser intensity for 1 second. Fluorescence recovery data was obtained immediately after bleaching and every 5 seconds up to 5 minutes.

### 2.5 Protein Purification and Nucleosome assembly

HP1α was purified as described previously^32^. Mono-nucleosomes with human histones were generated as described previously^33^. Histones used in this study are free of modifications.

Full length Ki-67 gene was synthesized (Twist Bioscience) and inserted into the yeast expression plasmid (-Ura) along with N-terminal hexa-histidine tag by Gibson assembly. Ki-67 was expressed in the S. cerevisiae strain BCY123. Starter cultures were grown to saturation overnight in CSM-Ura− media supplemented with 2 % dextrose, 2 % lactic acid, and 1.5 % glycerol at 30 °C. Starter cultures were then diluted 10-fold in YP media with 2 % lactic acid and 1.5 % glycerol, and grown to an OD of 1.0–1.3 at 30 °C (12–15 h), at which point protein expression was induced by the addition of 2 % galactose for 6 h at 30 °C. Cells were harvested by centrifugation, resuspended in 1 ml of 1 mM EDTA and 250 mM NaCl per liter of culture, and flash frozen dropwise in liquid nitrogen for storage at −80 °C.

For purification, frozen pellets were lysed by cryogenic grinding in a Freezer Mill (SPEX SamplePrep). The cell powder was resuspended in K-Buffer (50 mM Hepes-KOH (pH 7.5), 300 mM NaCl, 30 mM imidazole (pH 8.0), 10 % glycerol, 1 mM PMSF, 2.34 μM leupeptin, 1.45 μM pepstatin, and 0.5 mM TCEP). Lysate was clarified by centrifugation at 16,000 rpm in a JA 25.50 rotor for 45 min and loaded onto a HisTrap HP (GE) nickel-chelating Sepharose column. Protein was eluted in K-Buffer with 250mM NaCl and 500 mM Imidazole. Ki-67 fractions were loaded onto Hitrap-SP column and eluted with K-buffer containing 1M NaCl. Fractions containing Ki-67 were concentrated and loaded onto Superose6 10/300gl column pre-equilibrated in Storage buffer (50 mM Hepes-KOH (pH 7.5), 300 mM NaCl, 10% glycerol, 1 mM TCEP). Fractions containing Ki-67 were pooled, concentrated and stored in −80C.

### 2.6 DNA synthesis

DNA was synthesized from a Widom 601 sequence containing plasmid pGEMz_601 (Addgene, 26656) with primers containing a biotin and Cy3. AT-rich, GC-rich, and all primers were constructed by IDT:

125bp AT-rich DNA 5’AGCGGTGATGCTGATAGAAGTATAATATTAATAATAAATTAA ATATATTATATTAATAATTAATAATTAATAAATTAAAATATTA TTTATAATAATTAAACATAATAGCTTCTGTGCGCC – 3’
122bp GC-rich DNA 5’TGAACCTGTACCCTTGTTGGCGCGTACGCGCGAACGCGTTATCGTCGCGTACGCGCGACGCGACGCGCGATCGCGAACGCGCGTCGTCGCGCGACGCGCGGCCTTGTAGATGAACTTGCG – 3’
70bp DNA (−77N0) 5’CGTACGTGCGTTTAAGCGGTGCTAGAGCTGTCTACGACCAATTGAGCGGCCTCGGCACCGGGATTCTCCA – 3’
147bp DNA (0N0) 5’CAGGATGTATATATCTGACACGTGCCTGGAGACTAGGGAGTAATCCCCTTGGCGGTTAAAACGCGGGGGACAGCGCGTACGTGCGTTTAAGCGGTGCTAGAGCTGTCTACGACCAATTGAGCGGCCTCGGCACCGGGATTCTCCA – 3’
231bp DNA (43N43) 5’ACTATCCGACTGGCACCGGCAAGGTCGCTGTTCAATACATGCACAGGATGTATATATCTGACACGTGCCTGGAGACTAGGGAGTAATCCCCTTGGCGGTTAAAACGCGGGGGACAGCGCGTACGTGCGTTTAAGCGGTGCTAGAGCTGTCTACGACCAATTGAGCGGCCTCGGCACCGGGATTCTCCAGGGCGGCCGCGTATAGGGTCCATCACATAAGGGATGAACTCGG – 3’.

## 3. RESULTS AND DISCUSSION

### 3.1 SLBs provide a platform for measuring fluid properties and condensates at a single molecule level

Supported lipid bilayers (SLBs) are a widely used platform that mimics the cellular membrane. SLBs can be integrated into many surface-based techniques and allow for investigating fundamental membrane biology^25, 26, 30^. Recent work has shown that SLBs are a suitable platform for investigating membrane bound liquid-liquid phase separation (LLPS), highlighting how traditional LLPS studies such as circular droplet formation and fusion can be replicated in two dimensions^34^.

As a proof of concept, we first formed SLBs containing 94% POPC, 1% 18:1 rhodamine PE for SLB visualization, and 5% phosphatidylethanolamine mimic containing a biotinylated head group, 18:1 Biotinyl PE, and bound a biotin-labeled double-stranded DNA to the SLB through an avidin linkage. After 43N43 DNA (N denotes the 147bp 601 Widom positioning sequence with 43 bp linker DNA) was bound on the SLB, widefield imaging was performed to confirm the fluorescence homogeneity across the surface of the SLB (Figure S1A). This was directly compared to SLBs containing 1% 18:1 rhodamine PE (99% POPC). A small area was selectively photobleached and fluorescence recovery was observed with a halftime of 65 s for the DNA-bound SLB and 69 s for the SLB with no bound DNA (Figure S1B). From FRAP curves generated by the fluorescence recovery we determined the bulk diffusion coefficient of 0.59 ± 0.046 μm^2^/s for 18:1 rhodamine PE containing SLBs and 0.55 ± 0.044 μm^2^/s for 18:1 rhodamine PE-containing, DNA-bound SLBs. This indicates that the binding of DNA has little interference on the diffusion of lipids in SLBs. These values are also consistent with the diffusion coefficients of SLBs with membrane bound molecules determined in other studies ^21, 52, 53^.

By reducing the concentration of fluorescent molecules bound to the SLB, we could observe diffusion of single anchored DNA across the SLB surface (Fig 1A). Particle density is tuned to allow for individual particle tracking while maintaining high enough density to allow for formation of small condensates (Fig 1B, 1C). To test the efficacy of SLBs as a platform for observing condensation, we first bound a 230-base pair (bp) long DNA labeled with Cy3 to SLBs. Lateral movement of particles were captured with a diverse population of diffusion coefficients (Fig 1B, 1E). We then added 100 mM spermine, a 4+ charged polyamine. Previously, spermine has been shown to bridge adjacent DNA duplexes^35^ to promote condensate formation. After the addition of spermine, individual particles formed into brighter small condensates (or clusters) (Fig 1C, 1E). In addition to the immobile condensates, we also observed mobile particles as well as arrested individual particles. The population of mobile particles is likely dependent on the concentration as well as the type of condensing agent: charge-based condensation as in the case of spermine leads to charge inversion-mediated condensate dispersal at sufficiently high concentrations. Condensate formation on SLB-bound particles relies on the lateral movement of phospholipids within the bilayer. Thus, irreversible membrane deformation or destruction may interfere with lateral movement, giving a false impression of condensation. To rule out this possibility, we induced charge inversion by increasing the concentration of spermine to 500 mM, where the negatively charged DNA backbones become coated in positive charges from spermine. The resulting charge inversion is observed as dispersal of condensates formed by spermine (Fig 1D, Video1). Moreover, mobility of the surface bound particles recovers to levels before the addition of spermine (Fig 1D, 1E).

**Figure 1.**
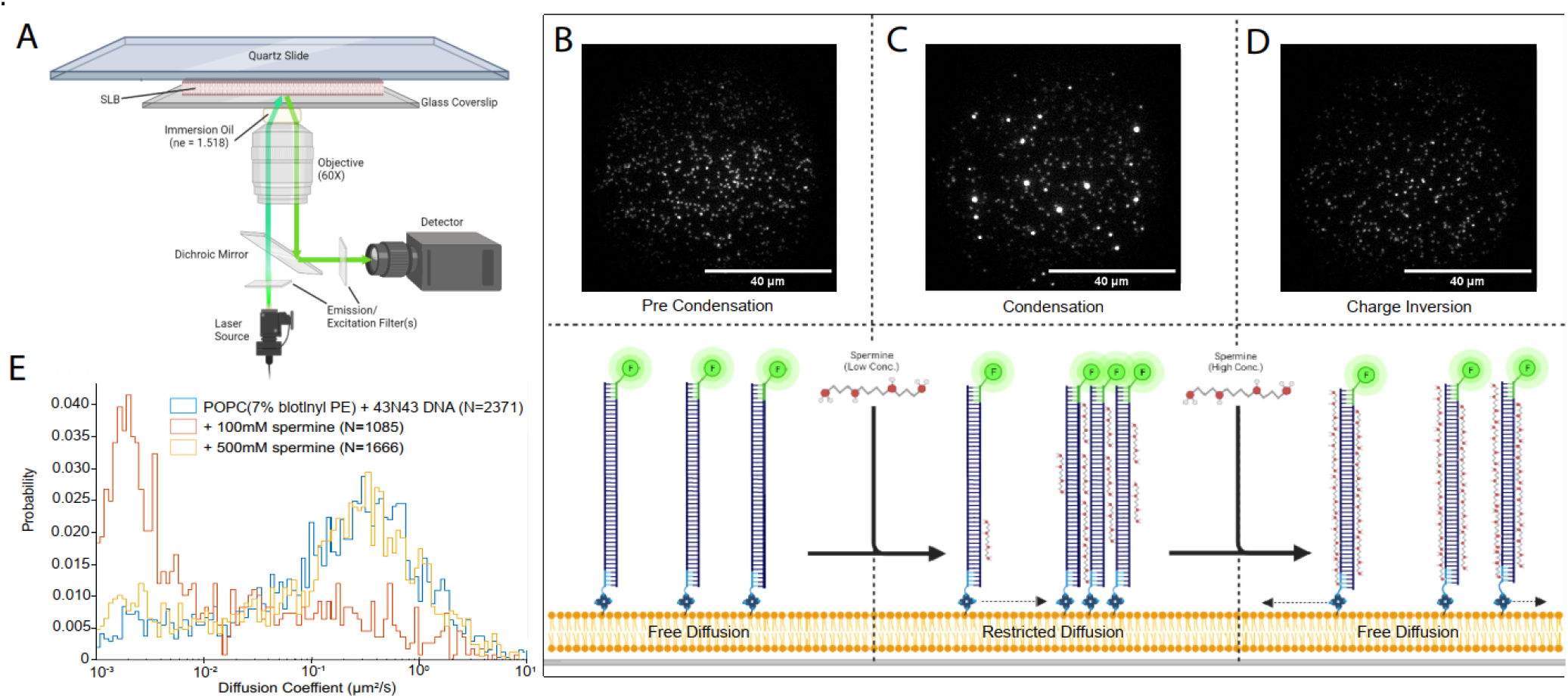
Experimental setup for experiments using objective-based total internal reflection fluorescence (oTIRF) microscopy in tandem with SLBs to observe spermine mediated condensation and dissolution: (A) oTIRF microscope setup, (B) visual representation and oTIRF recording of DNA bound to a 7% biotin containing SLB before the addition of any condensing agent, (C) visual representation and oTIRF recording of DNA bound to a 7% biotin containing SLB after promoting condensate formation through the addition of 100 mM spermine, (D) visual representation and oTIRF recording of DNA bound to a 7% biotin containing SLB after inducing charge inversion and condensate dispersal through the addition of 500 mM spermine, (E) diffusion histogram showing before, during, and after spermine mediated condensation and charge inversion of 40N40 DNA.

In this study, ‘initial condensation’ is described as the formation of bright, immobile puncta early when many particles are still mobile, and ‘definitive condensation’ is defined by a near complete loss of particle mobility and the ubiquitous formation of bright, immobile puncta (Fig 2A). We then explored how GC-content of DNA affects condensation. AT-rich DNA initially condensed after the introduction of 5 μM spermine, followed by 20 μM spermine for DNA with 57% GC content, and 30 μM for GC-rich DNA (Table 1). The individual concentration of condensation in each DNA sample is determined by titrating in 5 μM increments until condensates form. No initial condensation is observed below the concentration shown in Table1.

**Figure 2.**
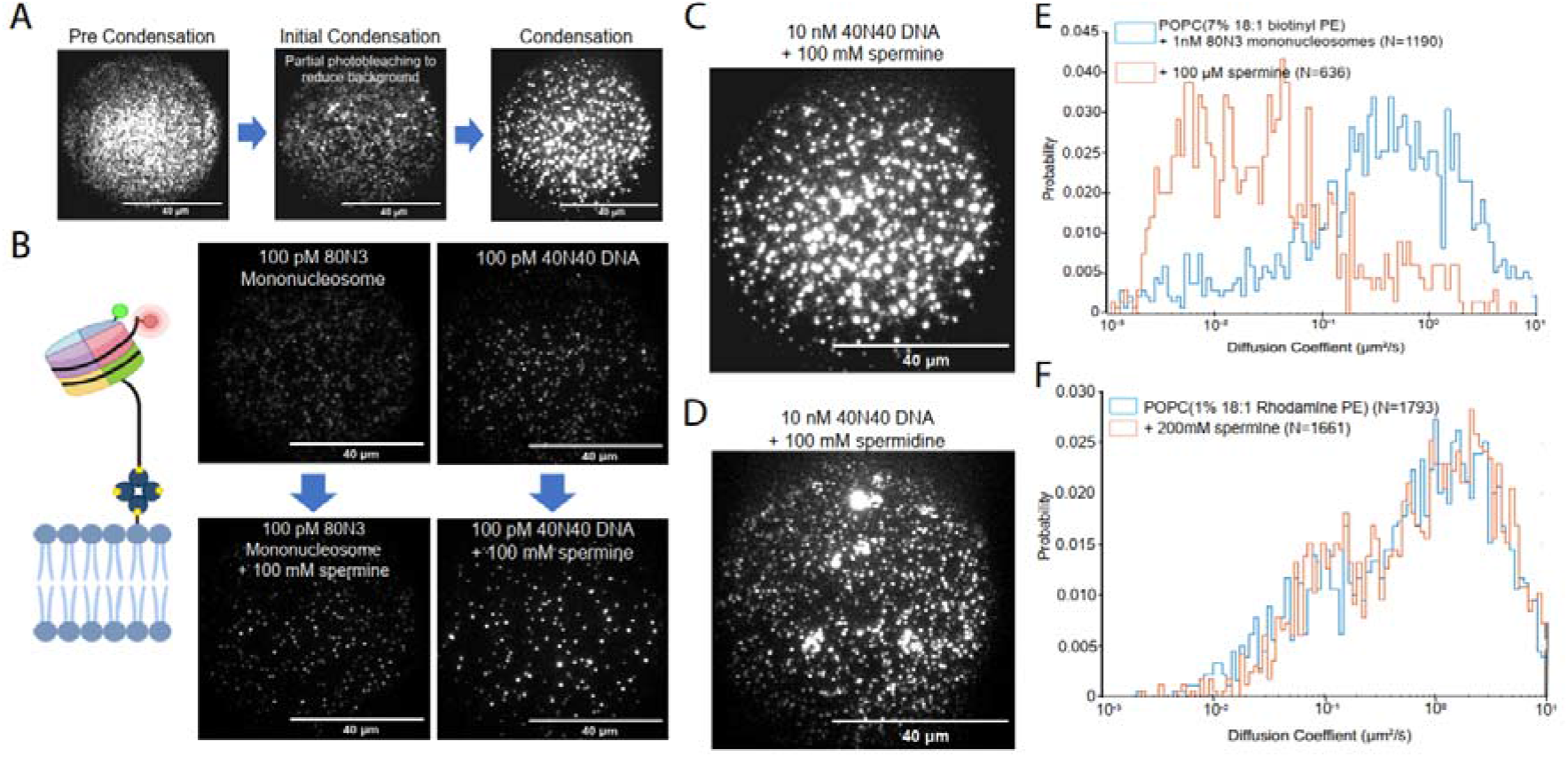
Condensation landscape: (A) definitions associated with condensation as they’re used in downstream experiments, (B) 80N3 mononucleosome construct used in subsequent experiments, example recordings of 80N3 nucleosomes and 40N40 DNA before and after condensate formation by spermine. (C) SLB populated densely with DNA in the presence of spermine. (D) SLB populated densely with DNA in the presence of spermidine. (E) diffusion histogram showing before, and after spermine mediated condensation of 80N3 nucleosomes, (F) diffusion histogram of a noncondensing particle (Rhodamine conjugated lipid) in an SLB.

**Table 1.**
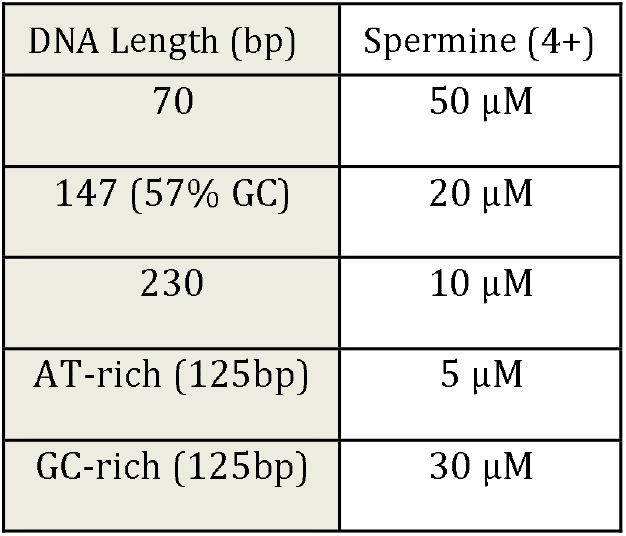
Condensability of different DNA lengths and GC-content by spermine.

The increased spermine-induced condensability of AT-rich DNA relative to GC-rich DNA confirms what was predicted from all-atom molecular dynamics simulations^35^. And the result is also consistent with frequency of transient contacts between two double stranded DNA molecules previous quantified by single-molecule FRET (smFRET)^35^. As we increased the length of DNA: from 70 bp 230 bp the threshold concentration of spermine for DNA condensation decreased from 50 μM to 10 μM (Table 1), indicating that longer DNA molecules are easier to condense through intermolecular interactions mediated by spermine.

To further expand this study toward condensation in the chromatin context, we generated a mono-nucleosome construct containing a 147 bp 601 nucleosome positioning sequence (denoted as N) flanked by an 80 bp linker containing biotin and a 3 bp linker containing Cy5 wrapped around a histone octamer with a Cy3 labeled H2A histone (Fig 2B). This ‘80N3’ nucleosome construct displays the same apparent two-dimensional diffusional behavior as naked DNA of the same length, and it can also form bright, immobile condensates upon addition of 100 mM spermine (Fig 2B). Using both 230 bp long DNA and 80N3 nucleosomes, we then tested several known and suspected condensing agents including polyamines, HP1α, and Ki-67 and determined the concentrations required to initiate condensation (Fig 2A, Table 2). Spermine (4+ valency) and spermidine (3+ valency) were both previously described to drive DNA condensation via electrostatic interactions^43^. We found that these polyamines behaved similarly when condensing DNA and 80N3 (Fig 2B, Table 2); however, a lower concentration of spermine was required to initiate DNA condensation than nucleosomes while the opposite is true for spermidine (Table 2). Interestingly, spermidine induced large, non-circular DNA condensates, where spermine at the same concentration, 100 mM (Fig 2C, 2D), only makes circular condensates. We also used the +2 charged polyamine putrescine and found that this was not sufficient to drive DNA condensation on the SLB even up to 1 M. We also tested HP1α, which binds to H3K9 methylated histones and promotes the formation of heterochromatin via recruitment of remodelers or binding partners^3^. We found that a 5-fold higher concentration of HP1α was required to initially drive 80N3 nucleosome condensation than DNA condensation, which occurred at HP1α concentrations as low as 10 nM (Table 2). Definitive condensation by HP1α occurred at 150 nM for DNA and at 200 nM for 80N3 nucleosomes (Fig. S2). Since HP1α has a DNA binding motif^48^, our studies further suggest that improved DNA accessibility for binding by condensing agents leads to improved condensation. We also note that HP1α-mediated nucleosome condensation occurred even in the absence of H3K9 methylation. We did not observe condensate dissolution even at the highest concentrations of HP1α or Ki-67 tested (10 μM or 1 μM, respectively).

**Table 2.** Condensing agent concentrations required to initiate DNA or nucleosome condensation on a 2D platform.

|  | 43N43 DNA | 80N3 Nucleosome |
| --- | --- | --- |
| Spermine (4+) | 10 $\mu\text{M}$ | 20 $\mu\text{M}$ |
| Spermidine (3+) | 30 $\mu\text{M}$ | 20 $\mu\text{M}$ |
| Putrescine (2+) | >1M | >1M |
| HP1α | 10 nM | 50 nM |
| Ki-67 | 1 nM | 100 pM |

### 3.2 Macromolecular condensation events can be visualized at the single molecule level in real time

Our SLB single molecule imaging platform also allows us to visualize DNA and nucleosome condensation in real time. Single particles were tracked to determine their mean-squared displacement (MSD) and subsequently, their diffusion coefficients. Cy3-labeled dsDNA bound to the SLB surface had an average diffusion coefficient of 0.49 ± 0.019 μm^2^/s with a lower limit of 0.001 μm^2^/s (Fig 1E). Single particle diffusion coefficients were measured before and after the addition of spermine and a clear shift from a high-mobility state to a low or no-mobility state was observed, from an average of 0.49 ± 0.019 μm^2^/s to 0.17 ± 0.019 μm^2^/s, with a shift back to 0.48 ± 0.025 μm^2^/s after the dispersal caused by spermine-mediated charge inversion (Fig 1E). A similar shift in diffusion coefficients was seen when subjecting 80N3 nucleosomes to the same spermine-mediated condensation, from an average of 0.94 ± 0.05 μm^2^/s to 0.17 ± 0.019 μm^2^/s (Fig 2E). The condensate formation and dispersal pattern were not seen, however, when performing the same experiments on fluorescent particles that do not form condensates such as rhodamine-labeled phospholipids (Fig 2F). Real-time single molecule experiments were carried out to visualize condensate formation and dissolution as the condensing agents were added via flow (Fig 3A). Once the condensates were fully formed, 500 mM spermine was added to promote condensate dispersal (Fig 3C) where we can see particles leave condensates and interact with other particles from dispersing condensates (Video 1, Fig 3C orange trace) or other condensates themselves (Video 1, Fig 3C yellow trace). From these videos, we were able to track the individual particles as they entered condensates by labeling each particle and observing their trajectories. We can then count the number of particles in a condensate by measuring the stepwise increase in intensity (Fig 3B) although transient interactions among condensate-forming particles coupled with photobleaching make it challenging to determine the number of particles-per-condensate reliably using intensity alone and other metrics may be needed.

**Figure 3.**
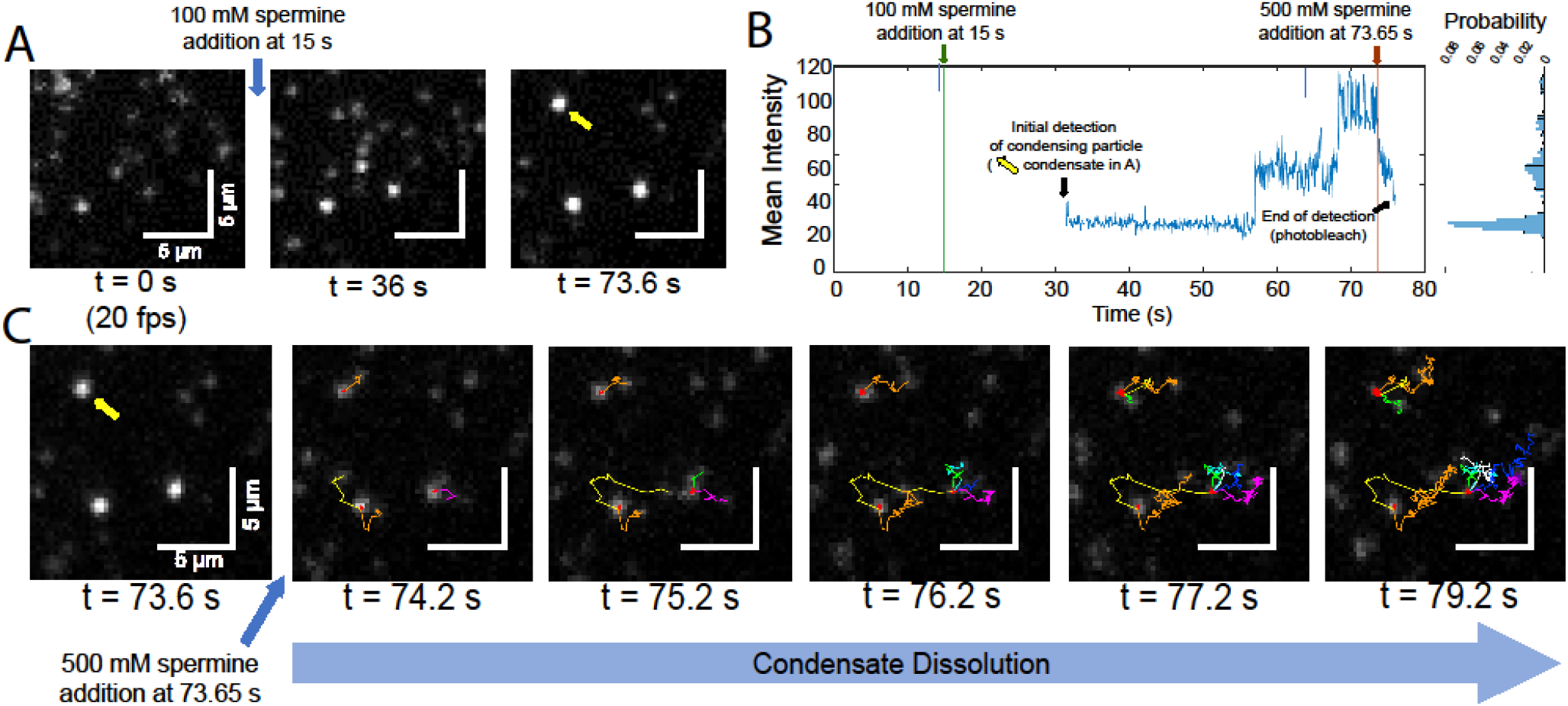
Realtime tracking of single molecules during condensation and dissolution: (A) snapshots highlighting condensate formation after the addition of 100 mM spermine. (B) Fluorescence intensity profile of a nucleating particle (denoted by a yellow arrow in A) during the inclusion of additional particles into the condensate. (C) Snapshots highlighting condensate dissolution after the addition of 500 mM spermine to the SLB in A. New particle trajectories are given a different color.

The large full width half maximum of the diffusion coefficient histograms seen in diffusing particles (Fig 1E, 2E blue histograms) indicates that the particles tend to have heterogeneous diffusion behaviors, which can also be visualized through direct observation (Video 2) and highlights the multiple diffusion modes observed within a population of SLB bound particles. To better track and categorize individual particles and their time-dependent diffusional behavior as they participate in condensation (Fig. 4A), we utilized a rolling MSD analysis in conjunction with a machine-learning algorithm termed diffusional fingerprinting^36^. From this analysis, we were able to categorize trajectories into subsegments of either a restricted or free movement state and determine which diffusional state a particle spends the most time in (Fig. 4B) and the specific diffusional metrics that differentiate these states (Fig 4C). In addition, we applied a fluorescence intensity analysis in order to observe a stepwise intensity increase during particle inclusion into condensates (Fig 3B) and during dissolution (Fig 4A). Interestingly, stepwise fluorescence intensity decreases can be seen when analyzing particles that have switched from restricted to free diffusion (Fig. 4A, middle and bottom panels) which is also observed prior to dissolution. This intensity decrease is likely due to particles leaving a condensate or through photobleaching. With this diffusional fingerprinting analysis, we ranked the key distinct trajectory metrics that discriminate restricted movement, seemingly characteristic of condensates, from free moving particles. Using a linear discriminate analysis dimensionality reduction, we found that the fractal dimension of trajectories was the most important metric distinguishing “Restricted” and “Free” diffusional states (Fig 4C). Trajectory step length distribution, the span of distances particles travel in a frame, and kurtosis, a measure of distribution ‘tailed-ness’, are also both important in identifying condensing particles (Fig 4C). In addition, five other metrics made contributions (Fig. 4C). When comparing diffusion characteristics of trajectories collected over the course of 60 seconds, 28 % of trajectories in the absence of spermine had more than one diffusion state vs 42% of trajectories in the presence of spermine (Table 3), likely due to particle inclusion into condensates.

**Figure 4.**
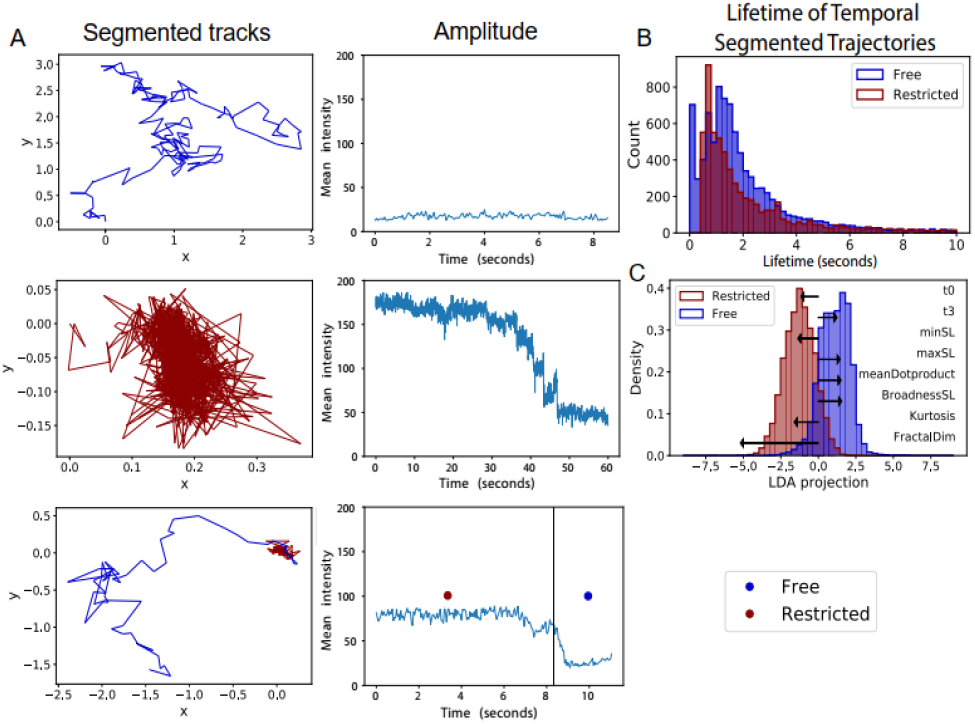
Diffusional fingerprinting of particles before and during dissolution: (A) Temporally segmented traces by rolling MSD analysis with their respective intensity profile. Showing a representative free moving trace (top), restricted trace (middle) and a heterogenous diffusing trace. Intensity profiles show that the free moving tracks have a generally constant intensity amplitude while the restricted trajectory has an intensity profile indicative of condensate disassembly. (B) The lifetime of free and restricted segments. (C) Shows a linear discriminate analysis dimensionality reduced representation of the diffusional fingerprints of free and restricted trajectory segments with the top 8 diffusional metrics separating the two distributions shown as black arrows. T0 and t3 describe residence times in the state with highest and lowest step length respectively.

**Table 3:** Temporal diffusional characteristics of 43N43 DNA before inducing condensation, after condensation is induced, and when condensation and dissolution are initiated by spermine concurrently in the same SLB as in Figure 3. Obtained from diffusional fingerprinting analysis and rolling MSD analysis where the diffusional states refer to “restricted” and “free”.

|  | Particles with<br>>1<br>Diffusional<br>State | Particles with >2<br>Diffusional<br>States | Particles with >1<br>Diffusional State<br>that end restrict-<br>ed | Particles that<br>are restricted<br>their entire<br>lifetime |
| --- | --- | --- | --- | --- |
| Pre-condensation | 28 % | 14 % | 15 % | 16 % |
| Post-<br>condensation | 42 % | 26 % | 23 % | 29 % |

### 3.3 Ki-67 driven nucleosome condensation

We determined that Ki-67 is capable of condensing 1 nM mononucleosomes incubated on an SLB at as low as 300 pM Ki-67 concentration (Fig 5A). To investigate particle dynamics within Ki-67 condensates, we performed two-color experiments using 0.25 nM Cy5 labelled 80N3 nucleosomes and 100 nM Cy3 labelled 40N40 nucleosomes. This scheme allowed for the formation of large condensates that can be observed with Cy3 at low laser intensity while the low concentration of Cy5 labelled nucleosomes allows for the tracking of single particles within and around the Cy3-verified condensates (Fig 5B). Borders of these Cy3-verified condensates are defined by a minimum fluorescence intensity of 90 (arbitrary unit) or triple the intensity of a single particle (Fig 5B, yellow borders). Colocalized with these large condensates is a significant proportion of particles that are stationary as well as dense immobile condensates seen both by Cy3 and by sparsely labelled Cy5 (Fig 5B, blue arrows) which implies that these large loosely packed condensates are composed of a network of the smaller, bright, dense condensates (Fig 2D, 5B) reminiscent of the dense bright condensates seen in previous experiments (Fig 2C). Visually, large condensates appear to take on a multitude of shapes but are immobile and seldom form circular condensates that are typically associated with phase-separated condensates. Nevertheless, the presence of particles capable of associating with, dissociating from, and travelling within or through a Ki-67 condensate indicates that these large condensates have partial-fluid like properties (Figure S3). Single particle trajectories that associate with large condensates are observed by Cy5 emission to undergo sudden diffusional changes, seemingly as a particle stops interacting with a condensate (Fig 5B, S3). This is further supported by stepwise fluorescence intensity changes seen concurrently with sudden changes in a particle’s MSD (Fig. 5B). These data suggest that, after inducing condensation for 5 minutes, while these large condensates have static position and shape, they do retain some fluid like properties associated with LLPS. Ki-67 condensates were not directly observed fusing to form larger condensates.

**Figure 5.**
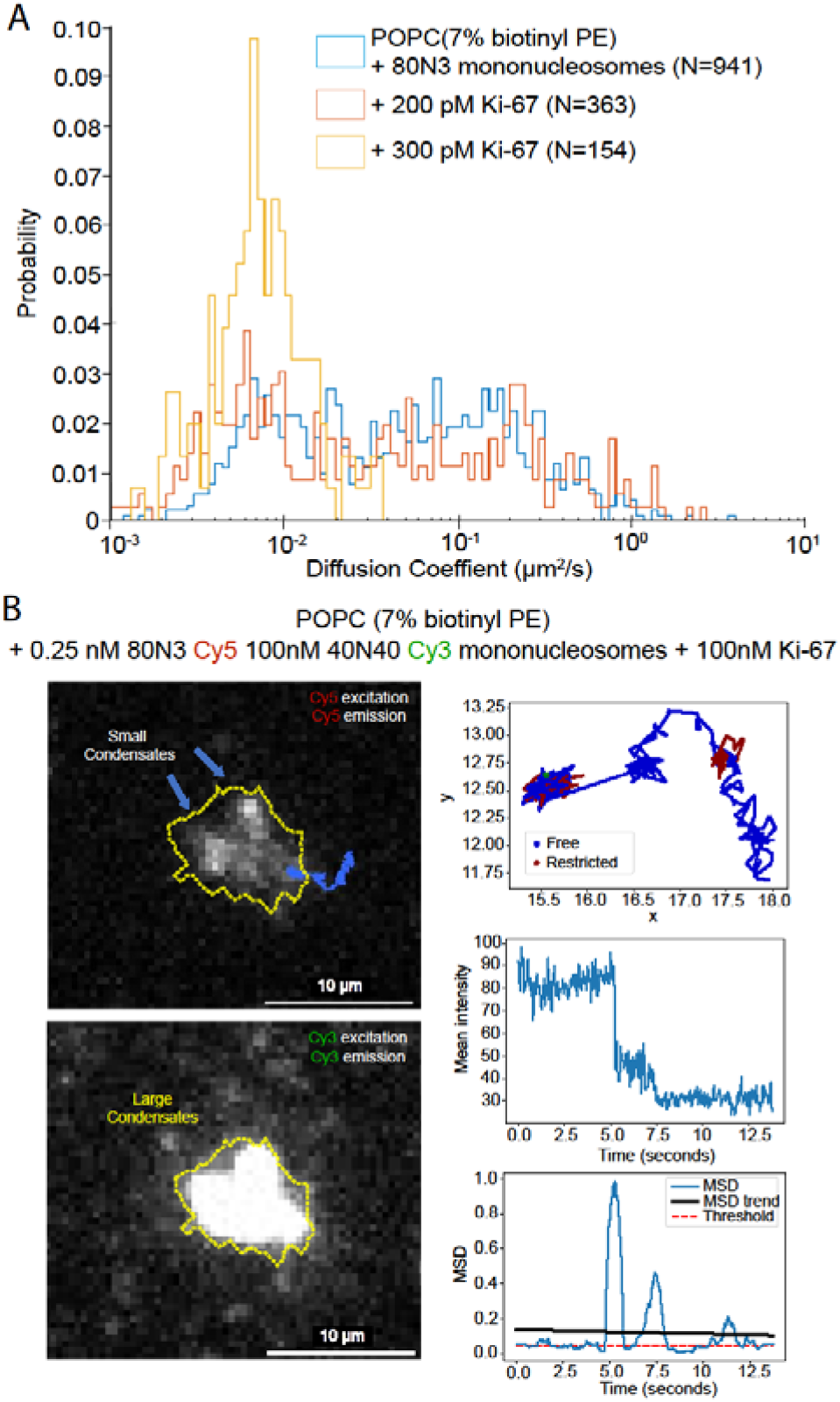
(A) Overlay of diffusion histograms obtained from tracked particles after the addition of 200 pM, and 300 pM Ki-67, (B) Large condensate sparsely labelled with Cy5 40N40 nucleosomes in a background of Cy3 labeled 80N3 nucleosomes. Condensate is outlined with a yellow dotted line and the trajectory of an interacting particle highlighted in blue; intermediate sized immobile condensates are shown using blue arrows (left panels). Trajectory specific diffusion/ fluorescence intensity/ rolling MSD profiles generated by diffusional fingerprinting (right panels).

## 4. CONCLUSIONS

In this work, we have used supported lipid bilayers as a platform for visualizing the 2-dimensional diffusion of surface-bound macromolecules such as DNA and nucleosomes in the presence of different condensing agents. We then observed that the threshold concentration of condensing agents depends on the length and GC content of the DNA, where DNA of greater length and lower GC contents led to initial condensation at lower condensing agent concentrations. Furthermore, DNA condensation by spermine can be reversed upon the addition of a DNA saturating concentration of spermine, highlighting that the condensation does not perturb the SLB and allows for the observation of condensation and dissolution as reversible processes. We then used SLBs for visualizing real time condensation and dispersal of condensates which allows tracking of individual particles entering and exiting condensates. These particle tracking data imply the presence of multiple diffusion states as indicated by the wide range of diffusion coefficients of DNA even in the absence of condensing agents. Using diffusional fingerprinting and temporal segmentation of diffusion we confirmed the existence of multiple diffusional states in DNA trajectories, namely a mobile and immobile state and identified their most discriminative diffusional characteristics.

We expanded this study by investigating the condensing behavior of nucleosomes in the presence of several known and suspected condensing agents such as Ki-67. Many proteins that participate in phase-separated condensates achieve condensation through multivalent interactions, where particles with a valency of 2 cannot form space-spanning interacting networks without linking to higher valence molecules^39^. With evidence that full length Ki-67 is capable of nucleosome condensation, it stands to reason that multiple motifs are likely responsible for Ki-67 condensation. To this end, future directions include testing the nucleosome condensability by individual Ki-67 motifs as well as investigating the impact of dephosphorylated Ki-67^37^ on nucleosome condensation.

Overall, this study serves to highlight the utility of SLBs as a tool for studying real-time kinetics of nucleosome condensation and condensate dispersal, which can be further expanded to investigate other systems such as signal transducing biomolecular condensates that form as a result of membrane receptors binding their ligands^40^.

## ASSOCIATED CONTENT

### Supporting Information

Widefield images depicting the fluorescence recovery after photobleaching (FRAP) of supported lipid bilayers as they are either bound by a macromolecule or not, oTIRF images highlighting the initial condensation of DNA and nucleosomes by HP1, and oTIRF/ particle tracking analyses of particles as they interact with condensates.

## AUTHOR INFORMATION

### Authors

**Nils Benning** - Department of Biology, Johns Hopkins University, 3400 N. Charles St. Baltimore, MD 21218-2683, USA

**Jacob Kæstel-Hansen** – Department of Chemistry and Nanoscience Centre, University of Copenhagen, Oster Farimasgade 5, DK-1353 Copenhagen K, Denmark

**Fahad Rashid** - Department of Biophysics and Biophysical Chemistry, Johns Hopkins University School of Medicine, 725 N. Wolfe St. Baltimore, MD 21205, USA

**Sangwoo Park** – Department of Biophysics and Biophysical Chemistry, Johns Hopkins University School of Medicine, 725 N. Wolfe St. Baltimore, MD 21205, USA

**Raquel Merino Urteaga** – Department of Biology, Johns Hopkins University, 3400 N. Charles St. Baltimore, MD 21218-2683, USA

**Ting-Wei Liao** – Department of Biophysics, Johns Hopkins University School of Medicine, 725 N. Wolfe St. Baltimore, MD 21205, USA

**Jingzhou Hao** – Department of Biophysics, Johns Hopkins University School of Medicine, 725 N. Wolfe St. Baltimore, MD 21205, USA

**James M. Berger** – Department of Biophysics and Biophysical Chemistry, Johns Hopkins University School of Medicine, 725 N. Wolfe St. Baltimore, MD 21205, USA

**Nikos S. Hatzakis** – Department of Chemistry and Nanoscience Centre, University of Copenhagen, Oster Farimasgade 5, DK-1353 Copenhagen K, Denmark, Denmark

-Novo Nordisk Foundation Centre for Protein Research, Faculty of Health and Medical Sciences, University of Copenhagen, Blegdamsvej 3B, DK-2200 Copenhagen N, Denmark

### Author Contributions

All authors have given approval to the final version of the manuscript.

## ACKNOWLEDGMENT

The authors thank Dr. Matthew Poyton and Claudia Carcamo for their initial instruction on the formation and handling of supported lipids bilayers. This work was supported by the National Institutes of Health (GM122569 to T.H), NSF EFMA (1933303), The Villum Foundation (BioNEC 18333 & Villum experiment grant 40801), and The Novo Nordisk Foundation (NNF14CC0001). The authors wish to thank The Histone Source at Colorado State University for generating the Xenopus histone octamers used in this study.

## ABBREVIATIONS

LLPS: liquid-liquid phase separation
SLB: supported lipid bilayer
SUV: small unilamellar vesicle
TIRM: total internal reflection microscopy
TIRF: total internal reflection fluorescence
MSD: mean squared displacement
LAP: linear assignment problem
LoG: Laplacian of Gaussian
FRAP: fluorescence recovery after photobleaching
FRET: fluorescence resonance energy transfer

## REFERENCES

[1] Nadège, M.; Carmona-Gutierrez, D.; Madeo, F. Polyamines in Aging and Disease. Aging 3 2011 no. 8 (n.d.): 716–32.

[2] Harami, G. M.; Kovács, Z.J.; Pancsa, R.; Pálinkás, J.; Baráth, V.; Tárnok, K.; Málnási-Csizmadia, A.; Kovács, M. Phase Separation by SsDNA Binding Protein Controlled via Protein−protein and Protein−DNA Interactions. Proceedings of the National Academy of Sciences 2020, 117, no. 42: 26206–17.

[3] Larson, A. G.; Elnatan, D.; Keenen, M.K; Trnka, M.J.; Johnston, J.B.; Burlingame, A.L.; Agard, D.A.; Redding, .S.; Narlikar, G.J.; Liquid Droplet Formation by HP1α Suggests a Role for Phase Separation in Heterochromatin. Nature 547 2017, no. 7662: 236–40.

[4] Feric, M.; Vaidya, N.; Harmon, T.S.; Mitrea, D. M.; Zhu, L.; Richardson, T.M.; Kriwacki, R.W.; Pappu, R.V.; Brangwynne, C.P. Coexisting Liquid Phases Underlie Nucleolar Subcompartments. Cell 2016, 165, no. 7 :1686–97.

[5] Wang, B.; Zhang, L.; Dai, T.; Qin, Z.; Lu, H.; Zhang, L.; Zhou, F.; Liquid–Liquid Phase Separation in Human Health and Diseases. Signal Transduction and Targeted Therapy 6 2021, no. 1: 290.

[6] Li, Y. R.; Oliver D. King, O.D; James Shorter, J.; Aaron D. Gitler, A.D.; Stress Granules as Crucibles of ALS Pathogenesis. Journal of Cell Biology 201 2013, no. 3: 361–72.

[7] Polymenidou, M.; Cleveland, D.W.;. The Seeds of Neurodegeneration: Prion-like Spreading in ALS. Cell 147 2011, no. 3: 498–508.

[8] Patel, A.; Lee, H.O.; Jawerth, L.; Maharana, S.; Jahnel, M.; Hein, M.Y.; Stoynov, S.; Mahamid, J.; Saha, S.; Franzmann, T.M.; et al. A Liquid-to-Solid Phase Transition of the ALS Protein FUS Accelerated by Disease Mutation. Cell 162 2012, no. 5: 1066–77.

[9] Schmidt, Hermann Broder, Ariana Barreau, and Rajat Rohatgi. “Phase Separation-Deficient TDP43 Remains Functional in Splicing.” Nature Communications 10, no. 1 (October 25, 2019): 4890.

[10] Ray, S.; Nitu Singh, N.; Rakesh Kumar, R.; Komal Patel, K.; Satyaprakash Pandey, S.; Debalina Datta, D.; Jaladhar Mahato, J.; Rajlaxmi Panigrahi, R.; Ambuja Navalkar, A.; Surabhi Mehra, A.; et al. α-Synuclein Aggregation Nucleates through Liquid–Liquid Phase Separation. Nature Chemistry 12 2020, no. 8: 705–16.

[11] Ambadipudi, S.; Biernat, J.; Riedel, D.; Mandelkow, M.;Zweckstetter, M. Liquid–Liquid Phase Separation of the Microtubule-Binding Repeats of the Alzheimer-Related Protein Tau. Nature Communications 8 2017, no. 1: 275.

[12] Zbinden, A.; Pérez-Berlanga, M.; De Rossi, P.; Polymenidou, M. Phase Separation and Neurodegenerative Diseases: A Disturbance in the Force. Developmental Cell 55 2020, no. 1: 45–68.

[13] Perdikari, T. M.; Murthy, A.C.; Ryan, V.H.; Watters, S.; Naik, M.T.; and Fawzi, N.L. SARS-CoV-2 Nucleocapsid Protein Phase-Separates with RNA and with Human HnRNPs. The EMBO Journal 39 2020, no. 24: e106478.

[14] Wang, J.; Shi, C.; Xu, Q.; Yin, H. SARS-CoV-2 Nucleocapsid Protein Undergoes Liquid–Liquid Phase Separation into Stress Granules through Its N-Terminal Intrinsically Disordered Region. Cell Discovery 7 2021, no. 1 : 5.

[15] Heinrich, B. S.; Maliga, Z.;; Stein, D. A.; Hyman A.A.; Whelan, S.P.J.; Palese, P. Phase Transitions Drive the Formation of Vesicular Stomatitis Virus Replication Compartments. MBio 9 2018, no. 5: e02290–17.

[16] Sanders, D.W.;Kedersha, N.; Lee, D.S.W.; Strom, A.R.; Drake, V.; Riback, J.A.; Bracha, D.; Eeftens, J.M.; Iwanicki, A.; Wang, A. et al. Competing Protein-RNA Interaction Networks Control Multiphase Intracellular Organization. Cell 181 2020, no. 2: 306–324.e28.

[17] Leslie, Mitch. Science. “Separation Anxiety”. https://www.science.org/content/article/sloppy-science-or-groundbreaking-idea-theory-how-cells-organize-contents-divides (Accessed April 12, 2022).

[18] Gitler, A.D.; James Shorter, J.; Taekjip Ha, T.; and Sua Myong, S. Just Took a DNA Test, Turns Out 100% Not That Phase. Molecular Cell 78 2020, no. 2: 193–94.

[19] Bracha, D.; Walls, M.T.; Wei, M.; Zhu, L.; Kurian, M.; Avalos, J.L.; Toettcher, J.E.;Brangwynne, C.P. Mapping Local and Global Liquid Phase Behavior in Living Cells Using Photo-Oligomerizable Seeds. Cell 175 2018, no. 6: 1467–1480.e13.

[20] Jan Akhunzada, M.; D’Autilia, F.; Chandramouli, B.; Bhattacharjee, N.; Catte, A.; Di Rienzo, R.; Cardarelli, F.; Brancato, G. Interplay between Lipid Lateral Diffusion, Dye Concentration and Membrane Permeability Unveiled by a Combined Spectroscopic and Computational Study of a Model Lipid Bilayer. Scientific Reports 2019,, no. 1: 1508.

[21] Koch, S.; Seinen, A.; Kamel, M.; Kuckla, D.; Monzel, C.; Kedrov, A.; Driessen, A.J.M Single-Molecule Analysis of Dynamics and Interactions of the SecYEG Translocon. The FEBS Journal 288, no. 7 : 2203–21.

[22] Kudalkar, E. M., Davis, T. N., & Asbury, C. L. Single-Molecule Total Internal Reflection Fluorescence Microscopy. Cold Spring Harbor protocols 2016, no. 5

[23] Johnson, J. M.;Ha, T.; Chu, S.; Boxer, S.G.; Early Steps of Supported Bilayer Formation Probed by Single Vesicle Fluorescence Assays. Biophysical Journal 83 2002, no. 6: 3371–79.

[24] Perez, T.D.; Nelson, W.J.; Boxer, S.G.; Kam, L. E-cadherin tethered to micropatterned supported lipid bilayers as a model for cell adhesion. Langmuir. 25 2005, no. 21 (December 6. 2005): 11963–11968.

[25] Chan, Y.M.; Boxer, S.G.. Model Membrane Systems and Their Applications. Model Systems/Biopolymers 11 2007, no. 6 (December 1, 2007): 581–87.

[26] Lengerich, B.; Rawle, R.J.; Boxer, S.G. Covalent Attachment of Lipid Vesicles to a Fluid-Supported Bilayer Allows Observation of DNA-Mediated Vesicle Interactions. Langmuir 26 2010, no. 11: 8666–72.

[27] Chung, M.;Boxer, S.G. Stability of DNA-Tethered Lipid Membranes with Mobile Tethers. Langmuir 27 2011, no. 9: 5492–97.

[28] Wei, M.; Elbaum-Garfinkle, S.; Holehouse, A.S.; Chen, C.C.; Feric, M.; Arnold, C.B.; Priestley, R.D.; Pappu, R.V.;Brangwynne, C.P. Phase Behaviour of Disordered Proteins Underlying Low Density and High Permeability of Liquid Organelles. Nature Chemistry 9 2017, no. 11: 1118–25.

[29] Kamagata, K.; Iwaki, N.; Hazra, M.K.; Kanbayashi, S.; Banerjee, T.; Chiba, R.; Sakomoto, S.; Gaudon, V.; Castaing, B.; Takahashi, H.; et al. Molecular Principles of Recruitment and Dynamics of Guest Proteins in Liquid Droplets. Scientific Reports 11 2021, no. 1: 19323.

[30] Grusky, D. S.; Moss, F.R.; Boxer, S.G. Recombination between 13C and 2H to Form Acetylide (13C22H–) Probes Nanoscale Interactions in Lipid Bilayers via Dynamic Secondary Ion Mass Spectrometry: Cholesterol and GM1 Clustering. Analytical Chemistry 94 2022, no. 27: 9750–57.

[31] Tinevez, J,; Perry, N.; Schindelin, J,; Hoopes, G.M.; Reynolds, G.D.; Laplantine, E.; Bednarek, S.Y.; Shorte, S.L.; Eliceiri, K.W. TrackMate: An Open and Extensible Platform for Single-Particle Tracking. Image Processing for Biologists 2017, 115: 80–90.

[32] Larson, A. G.; Elnatan, D.; Keenen, M.M.; Trnka, M.J.; Johnston, J.B.; Burlingame, A.L.; Agard, D.A.; Redding, S.; Narlikar, G.L. Liquid Droplet Formation by HP1α Suggests a Role for Phase Separation in Heterochromatin. Nature 547 2017, no. 7662: 236–40.

[33] Luger, K.; Rechsteiner, T.J., Richmond, T.J. Expression and Purification of Recombinant Histones and Nucleosome Reconstitution. In Chromatin Protocols edited by Peter B. Becker, 1–16. Totowa, NJ: Humana Press, 1999.

[34] Banjade, S.;Rosen, M.K. Phase Transitions of Multivalent Proteins Can Promote Clustering of Membrane Receptors. Edited by Anthony A Hyman. ELife 3 2014, e04123.

[35] Yoo, J.; Kim, H.; Aksimentiev, A.; Ha, T.; Direct Evidence for Sequence-Dependent Attraction between Double-Stranded DNA Controlled by Methylation. Nature Communications 7 2016, no. 1: 11045.

[36] Pinholt, H.D.; Bohr, S.R.N.;Iversen, J.F.; Boomsma, W.; Hatzakis, N.S. Single-Particle Diffusional Fingerprinting: A Machine-Learning Framework for Quantitative Analysis of Heterogeneous Diffusion. Proceedings of the National Academy of Sciences 118 2021, no. 31 : e2104624118.

[37] Sun, X.;Kaufman, P.D. Ki-67: More than a Proliferation Marker. Chromosoma 127 2018, no. 2: 175–86.

[38] Cuylen-Haering, S.; Petrovic, M.; Hernandez-Armendariz, A.; Schneider, M.W.G.; Samwer, M.; Blaukopf, C.; Holt, L.J.; Gerlich, D.W. Chromosome Clustering by Ki-67 Excludes Cytoplasm during Nuclear Assembly. Nature 587 2020, no. 7833: 285–90.

[39] Sanders, D.W.; Kedersha, N.; Lee, D.S.W.; Strom, A.R.; Drake, V.; Riback, J.A.; Bracha, D.; Eeftens, J.M.; Iwanicki, A.; Wang, A.et al. Competing Protein-RNA Interaction Networks Control Multiphase Intracellular Organization. Cell 2020,181, no. 2 : 306–324.e28.

[40] Jaqaman, K.; Ditlev, J.A.. Biomolecular Condensates in Membrane Receptor Signaling. Cell Signalling 2021, 69: 48–54.

[41] Strom, A.R.; Emelyanov, A.V.; Mir, M.; Fyodorov, D.V.; Darzacq, X.; Karpen, G.H. Phase Separation Drives Heterochromatin Domain Formation. Nature 2017, 547, no. 7662: 241–45.

[42] Hilbert, L.; Sato, Y.; Kuznetsova, K.; Bianucci, T.; Kimura, H.; Jülicher, F.; Honigmann, A.; Zaburdaev, V.; Vastenhouw, N.W. Transcription Organizes Euchromatin via Microphase Separation. Nature Communications 12 2021, no. 1: 1360.

[43] Strickfaden, H.; Tolsma, T.O.; Sharma, A.;Underhill, D.A.; Hansen, J.C.; Hendzel, M.J. Condensed Chromatin Behaves like a Solid on the Mesoscale In Vitro and in Living Cells. Cell 183 2020, no. 7: 1772–1784.e13.

[44] Rafiee, M.; Zagalak, J.A.; Sidorov, S.; Steinhauser, S.; Davey, K.; Ule, J.; Luscombe, N.M.; Chromatin-Contact Atlas Reveals Disorder-Mediated Protein Interactions and Moonlighting Chromatin-Associated RBPs. Nucleic Acids Research 49, no. 22: 13092–107.

[45] Zhang, L.; Geng, X.; Wang, F.; Tang, J.; Ichida, Y.; Sharma, A.; Jin, S.; Chen, M.; Tang, M.; Pozo, F.M. et al. 53BP1 Regulates Heterochromatin through Liquid Phase Separation. Nature Communications 13 2022, no. 1: 360.

[46] Quinodoz, S.A.; Jachowicz, J.W.; Bhat, P.; Ollikainen, N.; Banerjee, A.K.; Goronzy, I.N.; Blanco, M.R.; Chovanec, P.; Chow, A.; Markaki, Y.; et al. RNA Promotes the Formation of Spatial Compartments in the Nucleus. Cell 184 2021, no. 23: 5775–5790.e30.

[47] Childs, A. C.;Mehta, D.J.;Gerner, E.W. Polyamine-Dependent Gene Expression. Cellular and Molecular Life Sciences CMLS 60 2003, no. 7: 1394–1406.

[48] Kumar, A.;Kono, H.;. Heterochromatin Protein 1 (HP1): Interactions with Itself and Chromatin Components. Biophysical Reviews 12 2020, no. 2: 387–400.

[49] Cuylen, S.; Blaukopf, C.; Politi, A.Z.; Müller-Reichert, T.; Neumann, B.; Poser, I.; Ellenberg, J.; Hyman, A.A.; Gerlich, D.W. Ki-67 Acts as a Biological Surfactant to Disperse Mitotic Chromosomes. Nature 535 2016, no. 7611: 308–12.

[50] Raspaud, C.E.I.;Leforestier, A.;Livolant, F. Spermine-Induced Aggregation of DNA, Nucleosome, and Chromatin. Biophysical Journal 77 1999, no. 3: 1547–55.

[51] Keenen, M.M; Brown, D.; Brennan, L.D.; Renger, R.; Khoo, H.; Carlson, C. R.; Huang, B.; Grill, S.W.; Narlikar, G.J.; Redding, S. HP1 Proteins Compact DNA into Mechanically and Positionally Stable Phase Separated Domains. Edited by Sebastian Deindl and Kevin Struhl. ELife 10 2021, : e64563.

[52] Strom, A. R.; Biggs, R.J.; Banigan, E.J.; Wang, X.; Chiu, K.; Herman, C.; Collado, J.; Yue, Y.; Politz, J.C.R.; Tait, L.J. et al. HP1α Is a Chromatin Crosslinker That Controls Nuclear and Mitotic Chromosome Mechanics. Edited by Geeta J Narlikar, Kevin Struhl, and Sy Redding. ELife 10 2021, : e63972.

[53] Machida, S.; Takizawa, Y.; Ishimaru, M.; Sugita, Y.; Sekine, S.; Nakayama, J.; Wolf, M.; Kurumizaka, H. Structural Basis of Heterochromatin Formation by Human HP1. Molecular Cell 69 2018, no. 3: 385–397.e8.

