## Supporting Information for "Dimensional Reduction for Single Molecule Imaging of DNA and Nucleosome Condensation by Polyamines, HP1α and Ki-67"

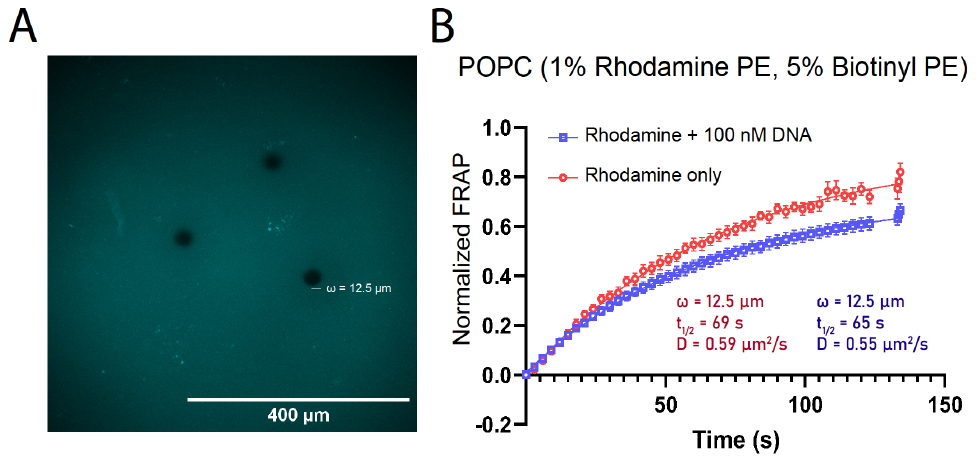

**Figure S1.** Bulk diffusion measurements of an SLB containing 1% of a dye labeled lipid and 5% biotinylated lipids: (A) widefield images of saturated SLBs during the bleaching step of a FRAP experiment on the bilayer (B) resulting fluorescence recovery curve after the bleaching seen in (A) and before and after the addition of 100 nM DNA. Insets provide data obtained from a one-phase association least squares fit.

S1

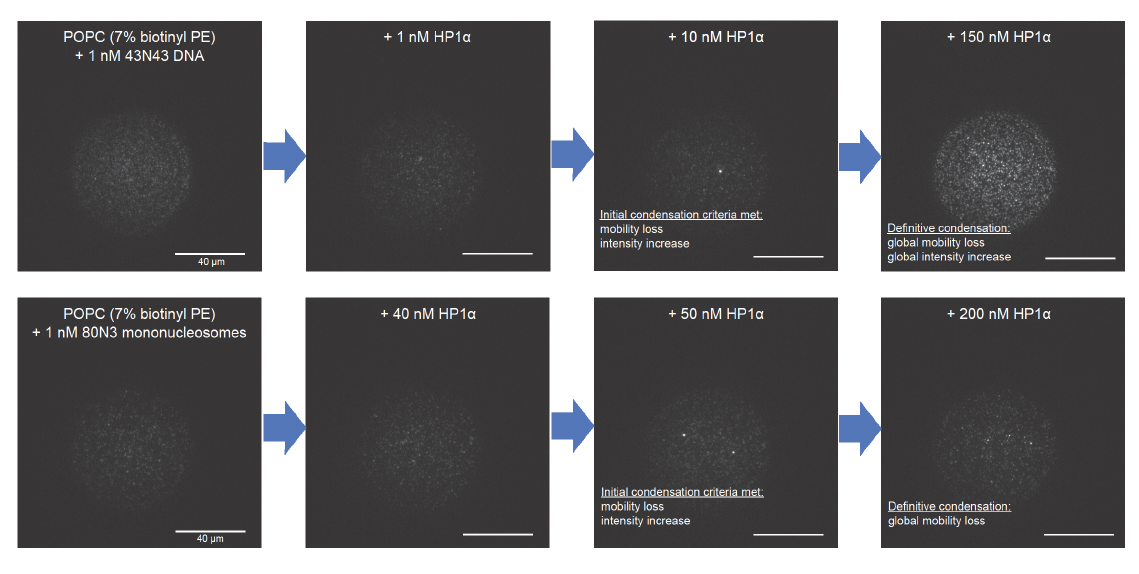

**Figure S2.** oTIRF images outlining the criteria for HP1α mediated DNA and nucleosome condensation. Criteria utilized to generate **Table 2.**

S2

**Figure S3.** Data highlighting the behavior of particles as they interact with condensates where columns represent one entire particle trajectory, data was collected in the same manner as **Figure 5,** with obstructed condensates outlined with a yellow dotted line. Each column corresponds to a trajectory seen interacting with a large condensate. When multiple trajectories are shown within the same condensate, the trajectory and its corresponding data are labelled with a number (1 through 4). Rows correspond to the same data type: (A) oTIRF image of Cy5 emission, (B) oTIRF image of Cy3 emission, (C) temporally segmented trajectory showing free and restricted movement of the particle, (D) fluorescence intensity of the particle over time, (E) rolling MSD and (F) a Pearson and Spearman test for correlation on fluorescence intensity vs MSD.

S3
